# Phospholipase B is critical for *Cryptococcus neoformans* survival in the central nervous system

**DOI:** 10.1101/2022.09.18.508401

**Authors:** Mohamed F. Hamed, Glauber Ribeiro de Sousa Araújo, Melissa E. Munzen, Marta Reguera-Gomez, Carly Epstein, Hiu Ham Lee, Susana Frases, Luis R. Martinez

**Author notes:** To whom correspondence may be addressed: Luis R. Martinez, University of Florida College of Dentistry Department of Oral Biology, 1395 Center Drive, DG-48, P.O. Box 100424 Gainesville, FL 32610.

## Abstract

*Cryptococcus neoformans* (*Cn*) is an opportunistic, encapsulated, yeast-like fungus that causes severe meningoencephalitis, especially in countries with high HIV prevalence. In addition to its well-known polysaccharide capsule, *Cn* has other virulence factors such as phospholipases, a heterogeneous group of enzymes that hydrolyze ester linkages in glycerophospholipids. Phospholipase B (PLB1) has been demonstrated to play a key role in *Cn* pathogenicity. In this study, we used a PLB1 mutant (*plb1*) and its reconstituted strain (Rec1) to assess the importance of this enzyme on *Cn* brain infection *in vivo* and *in vitro.* Mice infected with *plb1* strain survive significantly longer, have lower central nervous system (CNS) fungal load, and fewer and smaller cryptococcomas or biofilm-like brain lesions compared to H99- and Rec1-infected animals. *plb1* cryptococci are significantly more phagocytosed and killed by NR-9460 microglia-like cells. *plb1* cells have altered capsular polysaccharide biophysical properties that impair their ability to stimulate glia cell responses or morphological changes. We provide significant evidence demonstrating that *Cn* phospholipase is an important virulence factor for fungal colonization of and survival in the CNS as well as in the progression of cryptococcal meningitis. These findings may potentially help fill in a gap of knowledge in our understanding of cerebral cryptococcosis and may provide novel research avenues in *Cn* pathogenesis.

**IMPORTANCE:** Cryptococcal meningoencephalitis is a serious disease caused by infection of the neurotropic fungal pathogen *Cryptococcus neoformans* (*Cn*). Due to the increasing number of cases in HIV-infected individuals, as well as the limited therapies available, investigation into potential targets for new therapeutics has become critical. Phospholipase B (PLB1) is an enzyme synthesized by *Cn* that confers virulence to the fungus through capsular enlargement, immunomodulation, and intracellular replication. In this study, we examined the properties of PLB1 by comparing infection of *Cn* PLB1 mutant strain with both the wild-type and a PLB1 reconstituted strain. We show that PLB1 augments the survival and proliferation of the fungus in the CNS and strengthens virulence through modulation of the immune response and enhancement of specific biophysical properties of the fungus. The implications of PLB1 inhibition reveal its involvement in *Cn* infection and suggest that it may be a possible molecular target in the development of antifungal therapies. The results of this study support additional investigation into the mechanism of PLB1 to further understand the intricacies of *Cn* infection.

## INTRODUCTION

The encapsulated yeast-like fungus *Cryptococcus neoformans* (*Cn*) is an opportunistic pathogen that causes life-threatening meningoencephalitis in immunosuppressed individuals. Most cases of cryptococcosis are reported in Africa due to the high HIV infection rate and reduced access to standard-of-care including optimal antifungal drug therapy (1). The polysaccharide capsule is *Cn* main virulence factor. It is linked to the fungus ability to evade phagocytosis and suppress both cellular and humoral immunity (2, 3). However, there are other virulence factors that play a key role in *Cn* pathogenesis including the ability to grow at 37°C or mammalian temperature and the production of additional cell-associated factors such as melanin, which protects the fungus against environmental stress and antifungal drugs. Moreover, *Cn* produces less investigated degrading enzymes including proteinases (4), metalloprotease (5), DNase (6), urease (7), antioxidant (8), and lipases (9).

Phospholipases are classically classified into four major types, phospholipase A phospholipase B (PLB1), phospholipase C, and phospholipase D, based on the cleavage of ester linkages within a phospholipid molecule (10). *Cn* synthesizes a highly active extracellular phospholipase (11) with PLB1, lysophospholipase, and lysophospholipase-transacylase activities (9). Production of PLB1 in fungi correlates with virulence in mice (12) and decreased human neutrophil viability (13). *Cn* PLB1 triggers capsule enlargement, inhibits phagocytosis by macrophages, and is required for intracellular replication (14, 15). Disruption of the *plb1* gene markedly reduces all three enzyme activities without affecting cryptococcal virulence phenotypes, demonstrating that secretory PLB1 is a virulence factor for *Cn* (16). PLB1 is conveniently located in the cell wall via glycosylphosphatidylinositol anchoring, which allows immediate release of the enzyme in response to changing environmental conditions (17). PLB1 activity is required for cryptococcal pulmonary infection and for systemic dissemination from the respiratory system via the lymph nodes and blood vessels (18). Nevertheless, PLB1 is not essential for *Cn* blood-brain barrier (BBB) transmigration and brain invasion in mice especially via the trojan horse mechanism or inside of mononuclear phagocytes (18). Despite of this observation in rodents, *Cn* PLB1 was shown to promote fungal transmigration across an *in vitro* BBB model. It activates the host cell GTP-binding Rho family protein, Rac1, which regulates actin cytoskeleton (19), thus, suggesting a possible implication of PLB1 in fungal brain invasion and central nervous system (CNS) disease.

Although PLB1 is not required for *Cn* CNS invasion in rodents, it is not completely clear the importance of this enzyme in CNS colonization and progression of cerebral cryptococcosis. We investigated the role of *Cn* PLB1 on CNS infection and pathogenesis. We compared *Cn* CNS infection by a PLB1 mutant (*plb1*), its reconstituted (Rec1), and parental H99 strains (16). We provide significant evidence demonstrating that *Cn* PLB1 is an important virulence factor for fungal colonization of and survival in the CNS as well as in the progression of cryptococcal meningitis. Our findings may potentially help in our understanding of cerebral cryptococcosis and may offer novel research avenues in the study of *Cn* pathogenesis.

## RESULTS

### *Cn plb1* strain-infected mice survive longer than mice infected with H99 or Rec1 strains

We investigated the importance of *Cn* PLB1 in cerebral cryptococcosis by comparing the virulence of H99, *plb1*, and Rec1 strains in C57BL/6 mice infected systemically (Fig. 1). Rodents infected with H99 (*P*<0.05; median survival 13-days post-infection; dpi) and Rec1 (*P*<0.05; median survival 15-dpi) strains showed significantly faster mortality than mice infected with *plb1* (median survival 19-dpi) strain (Fig. 1A). Although both H99- and Rec1-infected mice began dying 11-dpi, 100% mortality occurred 13-dpi for the wild-type and 16-dpi for the complemented strain. *plb1*-challenged animals started dying 14-dpi and all of them were dead by 21-dpi.

**Fig. 1.**
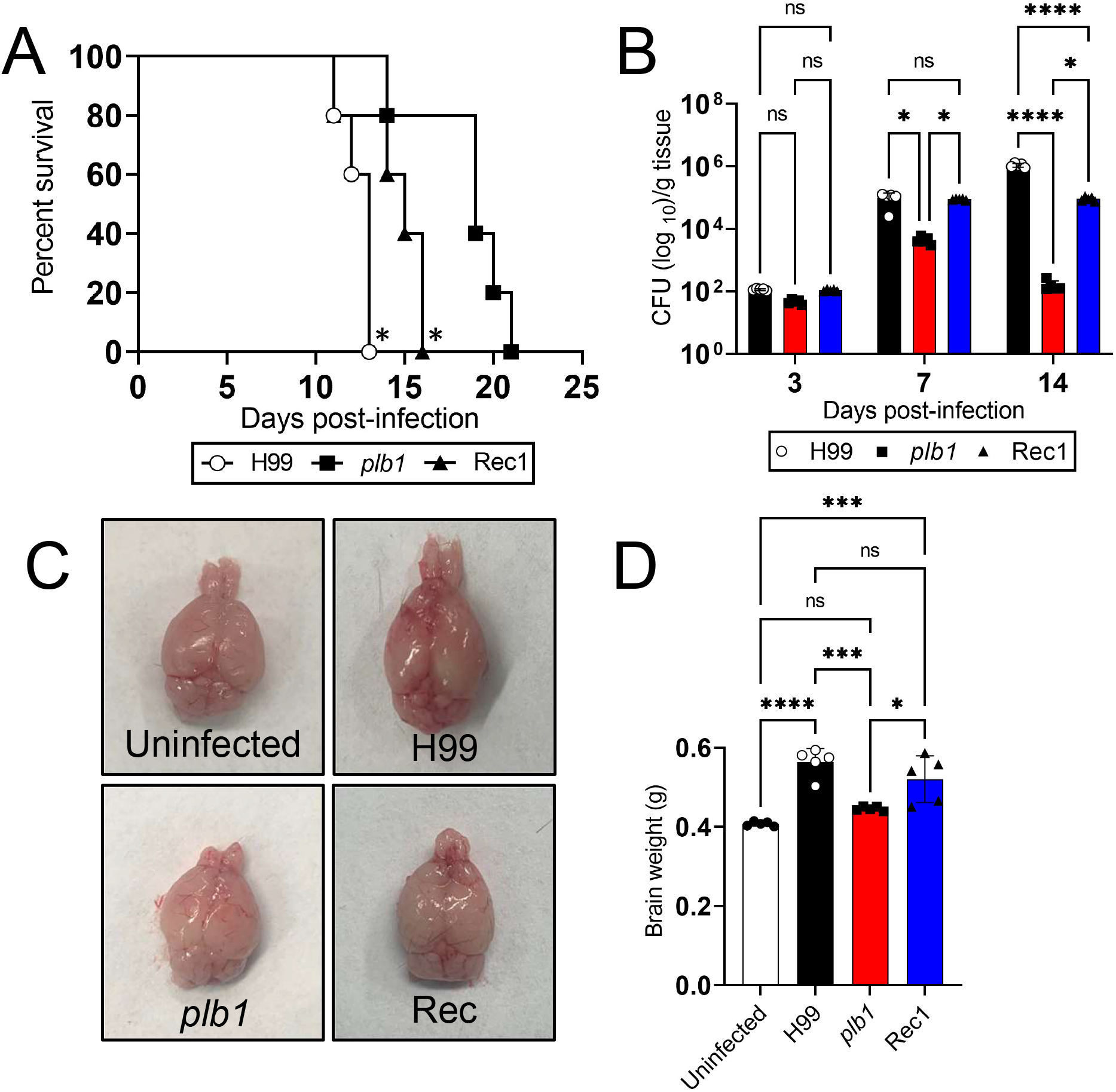
*C. neoformans* (*Cn*) phospholipase (PLB1) is critical for central nervous system (CNS) survival in systemically infected C57BL/6 mice. **(A)** Survival differences of C57BL/6 mice intravenously infected with 10^5^ *C. neoformans* strains H99, *plb1*, or Rec1 (*n* =7 per group). *P* value significance (*P*□<□ 0.05) was calculated by log rank (Mantel-Cox) analysis. Asterisk (*) denotes higher mortality than *plb1*-infected animals. **(B)** Fungal burdens (colony forming units; CFU) in brains collected from *Cn* H99-, *plb1*-, or Rec1-infected mice with 10^5^ cryptococci (*n* = 5 per group) at 3-, 7-, and 14-days post-infection (dpi). **(C)** Gross anatomy and **(D)** weight of brains (*n* =5 per group) excised from uninfected and infected C57BL/6 mice with H99, *plb1*, or Rec1 at 7-dpi. For **B** and **D**, bars and error bars denote the means and standard deviations (SDs), respectively. Asterisks denote *P* value significance (****, *P*□<□ 0.0001; ***, *P*□<□0.001; *, *P*□<□ 0.05) calculated by analysis of variance (ANOVA) and adjusted using Tukey’s post hoc analysis. ns denotes comparisons that are not statistically significant.

*Cn* PLB1 is not essential to cross the BBB and establish neurological disease in mice (18). However, the role of PLB1 in *Cn* survival is the CNS has not been investigated. We performed colony forming unit (CFU) determinations to understand the importance of *Cn* PLB1 on brain tissue survival (Fig. 1B). We confirmed that fungal PLB1 is not indispensable for brain invasion given that there were no differences in fungal burden in tissue homogenates from mice infected with H99 (1.15 × 10^2^ CFU/g tissue), *plb1* (4.64 × 10^1^ CFU/g tissue), or Rec1(1.09 × 10^2^ CFU/g tissue) 3-dpi. On day 7 post-infection, the fungal load in brain tissue increased for all the groups, although H99 (9.86 × 10^4^ CFU/g tissue; *P*<0.05)- and Rec1 (8.92 × 10^4^ CFU/g tissue; *P*<0.05)-infected mice displayed higher cryptococcal burden than those infected with *plb1* (4.4 × 10^3^ CFU/g tissue). On day 14 post-infection, brains excised from H99-infected animals showed the highest fungal load (1.1 × 10^6^ CFU/g tissue; *P*<0.0001 compared to the other groups) and followed in the order Rec1 (9.14 × 10^4^ CFU/g tissue; *P*<0.05 compared to *plb1*) > *plb1* (1.56 × 10^2^ CFU/g tissue). Interestingly, Rec1 CFU isolated from brain tissue of C57BL/6 mice were similar from 7- to 14-dpi whereas the *plb1* load was considerably reduced (~2 logs) during the same period.

Brain gross anatomy examinations revealed increased size and prefrontal cortex hemorrhage in both, H99- and Rec1-infected C57BL/6 mice 7-dpi, but blood outflow was more noticeable in rodents infected with the wild-type strain (Fig. 1C). Brains from *plb1*-infected mice evinced similar size than brains removed from naïve uninfected controls. To validate the gross inspections of the brain, weight measurements were performed (Fig. 1D), demonstrating that on average (*n* =7 per group), brains excised from mice infected with H99 (mean weight 0.564 g; *P*< 0.0001, *P*<0.001, and not significant (ns) relative to uninfected, *plb1*, and Rec1, respectively) and Rec1 (mean weight 0.521 g; *P*<0.001 and *P*<0.05 compared to uninfected and *plb1*, respectively) were heavier than those removed from uninfected (mean weight 0.408 g) and *plb1*-infected (mean weight 0.446 g) mice. There was no difference in brain weight between the uninfected and *plb1*-infected mice.

These results validated that *Cn* PLB1 is not crucial for CNS infection upon vascular dissemination, but it is required for fungal brain survival and persistence.

### *Cn plb1* cells form fewer and smaller cryptococcomas in the CNS of C57BL/6 mice

We investigated the impact of *Cn* PLB1 production and secretion in the CNS pathogenesis. For that, histopathological analyses of three affected regions of the mouse brain (e.g., cortex, hippocampus, and cerebellum) were performed 7-dpi with either *Cn* strain H99, *plb1*, or Rec1 (Fig. 2). In all neuroanatomical areas, each cryptococcal strain formed biofilm-like brain lesions or cryptococcomas, which were characterized by the presence of a central area of encephalomalacia or end-stage of liquefactive necrosis, loss of brain tissue, and minimal or no inflammation (Fig. 2A-C). Brain infection by H99 and Rec1 displayed encephalomalacia extended through the six laminar layers of the cerebral cortex whereas the *plb1*-infected cortex evinced a small circular area limited to laminar layers III and IV (Fig. 2A). Also, H99 and Rec1 formed higher number of cryptococcomas (*P*<0.01; *P*<0.001) and larger brain lesions (*P*<0.0001; *P*<0.0001) than *plb1* in cortical tissue (Fig. 2D). There were no differences in the number of cryptococcomas per cortical field between H99- and Rec1-infected mice (Fig. 2D).

**Fig. 2.**
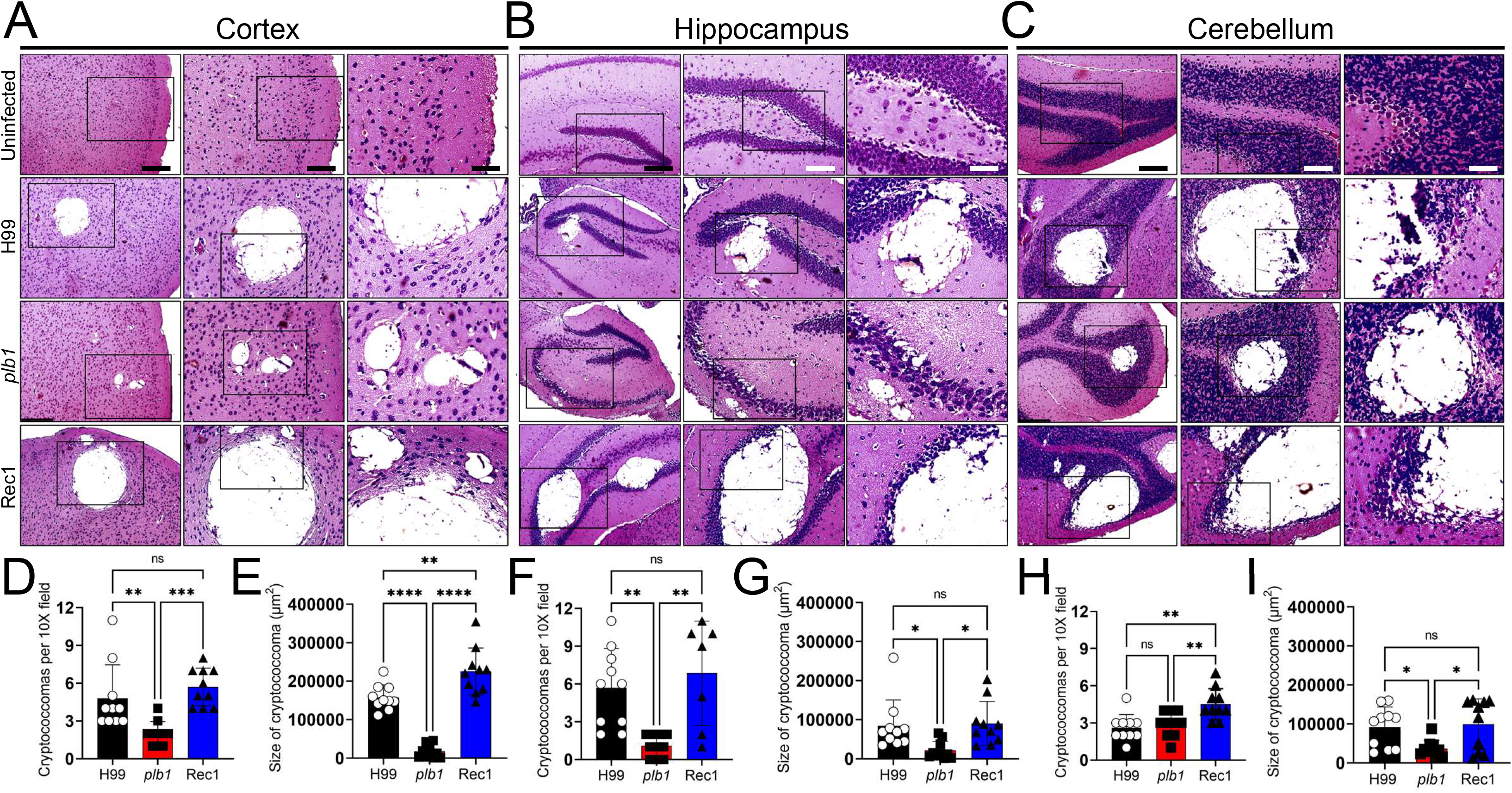
The brains of C57BL/6 mice systemically infected with phospholipase mutant (*plb1*) cryptococci exhibit reduced size and number or cryptococcomas (brain lesions). Histological examinations of the **(A)** cerebral cortex, **(B)** hippocampus, and **(C)** cerebellum from brains removed from *Cn* H99-, *plb1*-, or Rec1-infected mice with 10^5^ cryptococci at 7-dpi. Representative ×4 (left panel; scale bar, 200□μm), 10 (center panel; scale bar, 100□μm), and 20 (right panel; scale bar, 50□μm) magnifications of H&E-stained sections of the brain are shown. Black rectangular boxes delineate the area magnified (left to right panels). Count and size analyses of brain lesions caused by *Cn* H99-, *plb1*-, or Rec1 cells in **(D-E)** cortical, **(F-G)** hippocampal, and **(H-I)** cerebellar tissue sections in mice. Cryptococcoma counts per ×10 field were performed using an inverted microscope. The areas of 10 brain lesions per condition were measured using NIH ImageJ software. For **D** to **I**, bars and error bars denote the means and SDs, respectively. Each symbol (circles, squares, or triangles) represents an individual ×10 field or cryptococcoma area (*n* = 10 per group). Asterisks denote *P* value significance (****, *P*□<□ 0.0001; ***, *P*□<□ 0.001; **, *P*□<□ 0.01; *, *P*□<□ 0.05) calculated by ANOVA and adjusted using Tukey’s post hoc analysis. ns denotes comparisons that are not statistically significant.

The size of the cryptococcomas in the cortex of Rec1-infected mice was larger than those found in H99 (*P*<0.01; Fig. 2E). In the hippocampus, a region involved in learning and memory (Fig. 2B), *plb1*-infected brains exhibited a few and small cryptococcomas in the cornu ammonis 1 (CA1) region, whereas H99- and Rec1-infected brains had many cryptococcomas (*P*<0.01; H99 and Rec1 compared to mutant strain) and large size lesions (*P*<0.05; H99 and Rec1 compared to *plb1*) replacing great part of the dentate gyrus (Fig. 2F-G). H99- and Rec1-infected brains show no differences in the number (Fig. 2F) and size (Fig. 2G) of cryptococcomas formed in the hippocampus. We also analyzed cryptococcal infection of the cerebellum, a region in the CNS involved in motor function and coordination, and found a large area of encephalomalacia in H99- and Rec1-infected brains, which replaced the granular layer and extended to the Purkinje cell layer, which is the main cell in that region (Fig. 2C). In contrast, *plb1*-infected brains evinced smaller encephalomalacia area restricted to the granular layer of the cerebellum. Rec1 formed significantly more cryptococcomas per field in the cerebellum relative to H99 (*P*<0.01) and *plb1* (*P*<0.01) (Fig, 2H). H99 and *plb1* showed no differences in the number of cerebellar lesions per field. Nevertheless, the wild-type (*P*<0.05) and complemented (*P*<0.05) strains demonstrated substantially larger cerebellar lesions than the mutant strains (Fig. 2I). Our findings indicate that *Cn* PLB1 is important for cryptococcoma formation, which is imperative for fungal survival in the CNS.

### *Cn plb1* strain releases less GXM in the CNS than H99 or Rec1 strains

GXM production and release is required for *Cn* pathogenesis and to cause cryptococcal meningoencephalitis. GXM distribution was analyzed by immunohistochemistry (IHC) in the cerebral cortex, hippocampus, and cerebellum of infected mice 7-dpi (Fig. 3). *plb1* showed a diminished GXM release in tissue of every brain region analyzed. In this regard, H99- and Rec1-infected brains displayed extensive GXM secretion (brown staining) around the cortical cryptococcomas, involving a large tissue area including blood capillaries (Fig. 3A). In *plb1*-infected cortical tissue, GXM release surrounding the fungal brain lesions was reduced, circumscribed to the periphery of the cryptococcomas, and distant from the blood capillary walls (Fig. 3A). Analysis of GXM distribution in the cortex demonstrated that the Rec1 secreted significantly more CPS than H99 (*P*<0.01) and *plb1* (*P*<0.0001) (Fig. 3D). Additionally, H99 exhibited larger GXM distribution than *plb1* (*P*<0.05) in the cortex of infected mice. In the hippocampus, *plb1*-infected brains presented GXM localized in the CA1 region (Fig. 3B). However, GXM distribution in the hippocampus of H99-infected brains, spread further around the area of encephalomalacia in the dentate gyrus, whereas Rec1-infected hippocampus showed considerable GXM dispersal from the wall of the cryptococcoma to the parenchyma of the dentate gyrus and CA1 to CA3. Like the cortex, hippocampal GXM distribution was significantly more extensive in the Rec1-infected brain region compared to mice challenged with H99 (*P*<0.05) or *plb1* (*P*<0.0001) (Fig. 3E). *plb1*-infected hippocampal tissue evinced less GXM distribution than H99 (*P*<0.01). In the cerebellum, *plb1* showed little GXM release in the white matter or the granular layer surrounding the area of encephalomalacia caused by the cryptococcoma (Fig. 3C). H99- and Rec1-infected cerebella, however, exhibited greater GXM distribution around the area of encephalomalacia in the white matter and the granular, Purkinje cell layers of the gray matter (Fig. 3C). *plb1*-infected cerebella evinced a significant decreased in GXM tissue distribution compared to H99-(*P*<0.05) and Rec1 (*P*<0.01)-infected tissue (Fig. 3F). Cerebella infected with H99 or Rec1 show no difference in CPS spreading. Our results show that *plb1* releases less GXM in specific regions of the CNS than H99 or Rec1, suggesting that PLB1 production may correlate with the CPS production, having important implications for fungal survival in brain tissue.

**Fig. 3.**
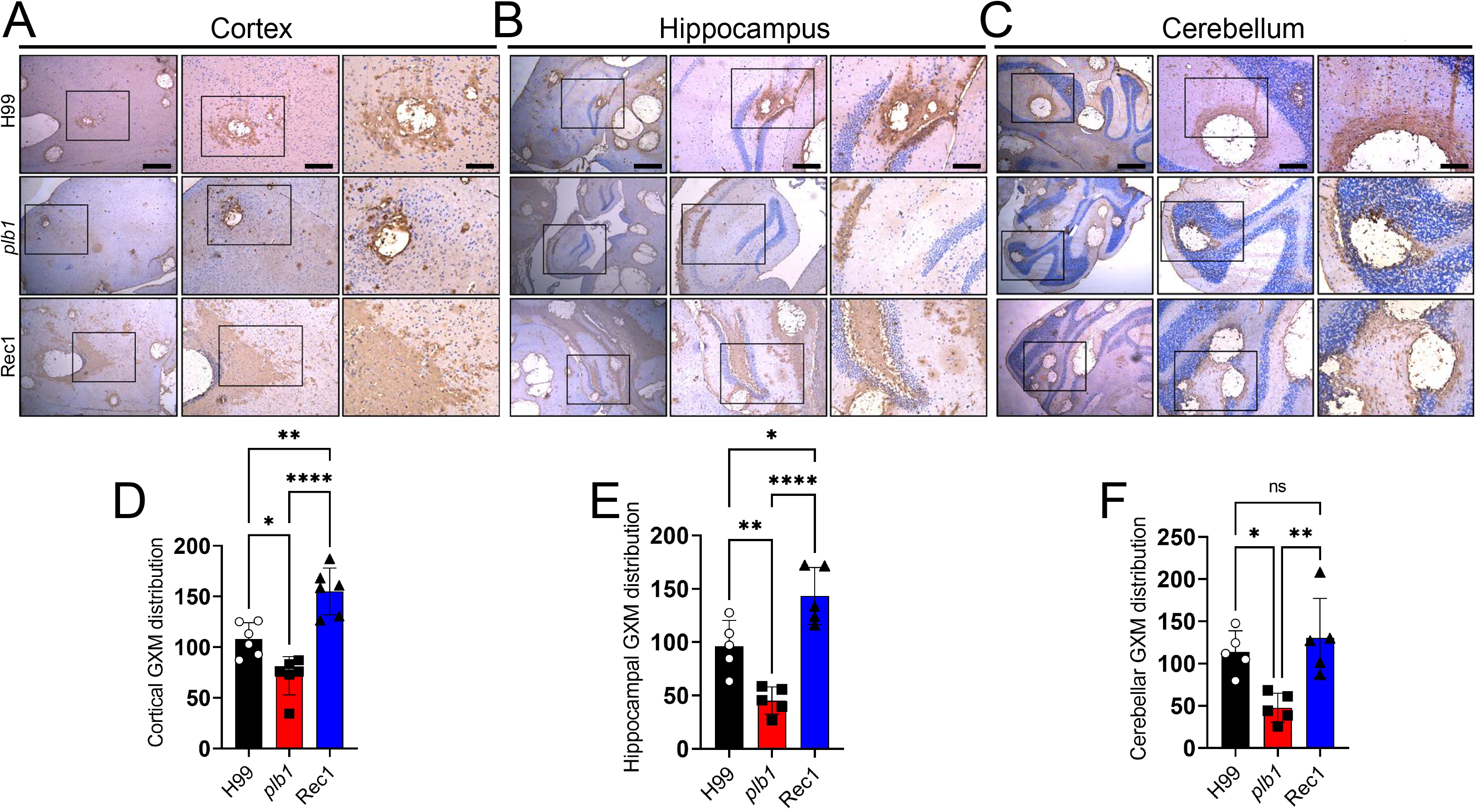
*Cn plb1* strain cells demonstrate reduced glucorunoxylomannan (GXM) release in brain tissue. Histological examinations of the **(A)** cerebral cortex, **(B)** hippocampus, and **(C)** cerebellum from brains removed from *Cn* H99-, *plb1*-, or Rec1-infected mice with 10^5^ cryptococci at 7-dpi. Representative ×4 (left panel; scale bar, 200□μm), 10 (center panel; scale bar, 100□μm), and 20 (right panel; scale bar, 50□μm) magnifications of GXM binding monoclonal antibody (mAb) 18B7-stained sections of the brain are shown. Black rectangular boxes delineate the area magnified (left to right panels). Brown staining indicates GXM secretion and accumulation. GXM distribution in **(D)** cortical, **(E)** hippocampal, and **(F)** cerebellar tissue sections. The areas of GXM distribution of 6 brain lesions per condition were measured using NIH ImageJ software. For **D** to **F**, bars and error bars denote the means and SDs, respectively. Each symbol (circles, squares, or triangles) represents an individual area (*n* = 6 per group). Asterisks denote *P* value significance (****, *P*□<□ 0.0001; **, *P*□<□ 0.01; *, *P*□<□ 0.05) calculated by ANOVA and adjusted using Tukey’s post hoc analysis. ns denotes comparisons that are not statistically significant.

### *Cn plb1* gene mutation results in biophysical changes to the CPS

*Cn* extensively releases its CPS and compromises the immune responses of the host to combat the infection (20). *plb1* had reduced CPS production around cryptococcomas. Given its importance in cryptococcal meningitis, we compared the biophysical properties of the secreted CPS by *Cn* strains H99, *plb1*, and Rec1 (Fig. 4). Scanning electron microscopy (SEM) images demonstrated that *plb1* and Rec1 yeast cells had similar cell sizes but smaller capsules and CPS fibers than H99 cryptococci (Fig. 4A). A Zeta potential analysis showed that *plb1* (−21.7±5.9 mV)-derived CPS evinced a higher reduction in its negative charge than H99 (−27.8±3.4 mV; *P*<0.05) and Rec1 (−27.7±4.5 mV; *P*<0.05; Fig. 4B). Remarkably, both genetically manipulated *Cn* strains, *plb1* (~135.3±1.3 µS; *P*<0.0001) and Rec1 (~144±1.4 µS; *P*<0.0001), had a significant decrease in CPS conductance compared to the parental strain H99 (~104.7±0.5 µS; Fig. 4C). We analyzed and compared the size distribution of the secreted CPS molecules from *Cn* H99, *plb1*, or Rec1 using dynamic light scattering (Fig. 4D). The CPS molecules from *plb1* yeast cells exhibited, on average, a smaller diameter (364.4±40 nm) than those of H99 (643.5±105.1 nm) and Rec1 (521±121 nm) cells (Fig. 4D). *plb1* yielded a widely distributed size population of CPS molecules (size range: 50.75-1943.09 nm; size average: 485.8 nm; size median: 129.38 nm). H99 and Rec1 produced two different size populations of CPS molecules. For H99, small size CPS molecules ranging 11.81 to 358.02 nm (size average: 121.47 nm; size median: 76.74 nm) and a large size ranging 1,548.87 to 1,587.70 nm (size average: 1,568.21 nm; size median: 1,568.16 nm). For Rec1, small size CPS molecules ranging 131.62 to 788.75 nm (size average: 344.50 nm; size median: 287.05 nm) and a large size ranging 1359.29 to 1,958.41 nm (size average: 1,642.68 nm; size median: 1,631.91 nm).

**Fig. 4.**
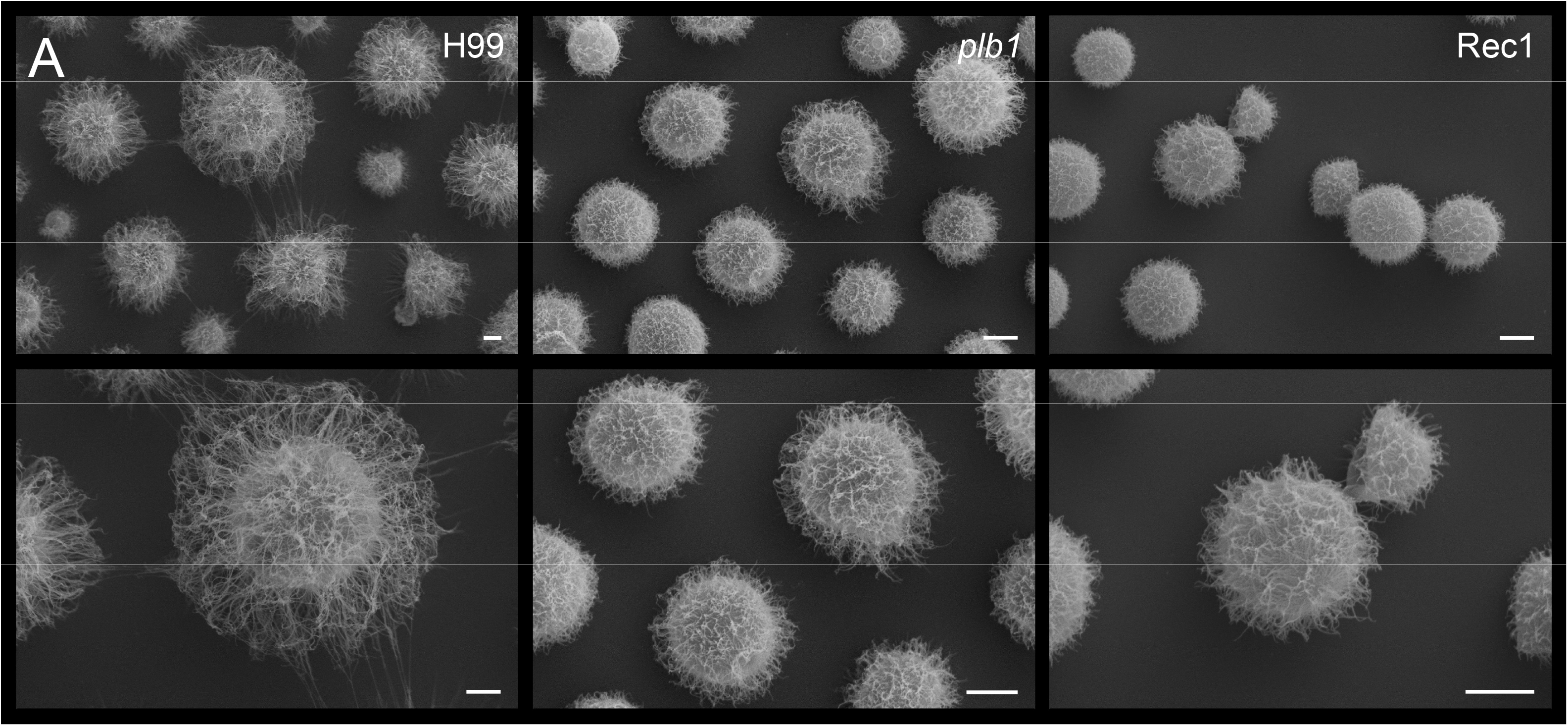
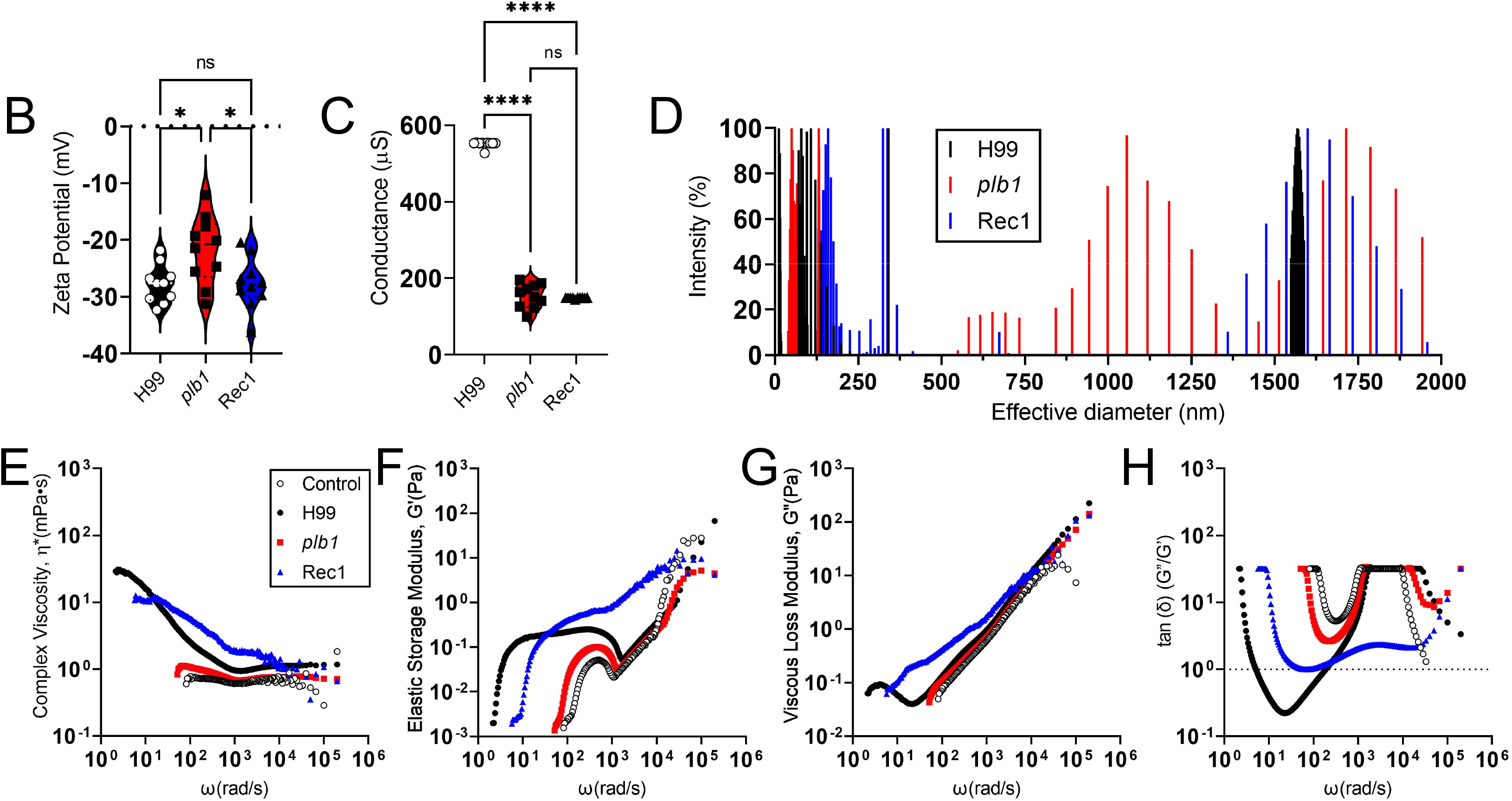
*Cn plb1* strain shows CPS alterations. **(A)** Scanning electron microscopy images of *Cn* H99, *plb1*, and Rec1, scale bar 2 μm. **(B)** Zeta potential and **(C)** conductance of secreted CPS by *Cn* strains H99, *plb1*, and Rec1. For **B** and **C**, each symbol represents the value for 1 fungal cell (*n* = 10 per group). Dashed lines represent averages of the results. Error bars denote SDs. Asterisks denote *P* value significance (*, *P* < 0.05; ns, not significant) calculated by ANOVA and adjusted using Tukey’s post hoc analysis. **(D)** CPS obtained from *Cn* strains H99, *plb1*, and Rec1. The *x* axis represents size distribution by particle diameter; the *y* axis corresponds to the values of percentage intensity-weighted sizes obtained with the nonnegative least-squares (NNLS) algorithm. The rheological properties of *Cn* strains H99, *plb1*, and Rec1 secreted polysaccharide were analyzed including **(E)** complex viscosity (□), **(F)** elastic storage modulus (G’), **(G)** viscous loss modulus (G”), and **(H)** viscoelastic behavior tan (δ) as a function of the stimulus angular frequency (ω). The dashed line in G indicates the value 1, where the behavior of the material changes.

Given of the different biophysical properties of the CPS secreted by H99, *plb1*, or Rec1, we studied their rheological behavior (Fig. 4E-H). For most fluids with long-chain microstructure such as polysaccharide fibers, the viscosity is a decreasing function of shear rate known as shear thinning (21–23). Complex materials respond to a force stimulus by storing energy (elastic behavior) or dissipating energy (loss behavior). Both behaviors are characterized by rheology techniques that allow the determination of the complex shear modulus, G*(ω) = G′(ω) iG′′(ω) where G′ is the rheological storage modulus and G′′ the rheological loss module, determined as a function of the angular frequency of the stimulus ω. In addition to the complex shear modulus, G*(ω) = G′ + iG′′, viscoelastic materials are also defined by their complex shear viscosity η ∗ (ω) = [G ∗ (ω)/(iω)] (21). We evaluated the viscosity of the η(ω), G′ (storage modulus) and G′′ (loss modulus) complex, measured at an angular stimulus frequency (ω) between 10 and 10^6^ rad/s. The complex viscosity curves in aqueous solution (10 mg/mL) at 37°C showed that, for all strains, the viscosity of the η(ω) complex decreased with increasing frequency (Fig. 4E). However, this effect is more pronounced in the *pbl1* mutant where it does not present stimuli below <10^2^ rad/s, having a similar behavior to the control (distilled-water). The storage module (G′) (Fig. 4F) and the loss module (G′′) (Fig. 4G) showed that they vary with frequency, although for H99 and Rec1 there is a spectral mechanical change at frequencies below 10^2^ rad/s, antagonistically, frequencies above <10^3^ rad/s G′ and G′′ converged to the maximum in all strains. The ‘damping behavior’ represented by tanδ (Fig. 4H) varied with the frequency, yet *plb1*, showed a reduction in the ability to stimuli.

We demonstrated that cryptococci with *plb1* gene mutation and to some extent its complemented strain Rec1 differ substantially in the structural composition and production of CPS molecules relative to H99 cells. These alterations of the CPS might have adverse implications on host-pathogen interactions and enhance CNS pathogenesis.

### *Cn* PLB1 mutant strain does not alter microglial morphology in cortical tissue

Microglia play a critical role in responding to eukaryotic pathogens by regulating inflammatory processes proficient at controlling CNS colonization. We assessed microglial responses and morphological changes in the cortex of C57BL/6 mice infected with *Cn* strains H99, *plb1*, or Rec1 7-dpi using IHC and light microscopy. Infiltrating microglia to the areas of fungal infection were stained with ionized calcium binding adaptor protein (Iba-1)-binding monoclonal antibody (mAb; Fig. 5), the most acceptable and used microglial marker. Uninfected mice showed the presence of naïve microglia characterized by a cylindrical soma or cell body surrounded by thin processes or ramifications/branches (Fig. 5A). H99- and Rec1-infected brains showed considerable number of microglia recruitment near the cryptococcomas with diverse morphology (Fig. 5A). Interestingly, *plb1*-infected tissue presented low number of ramified microglia proximal to the brain lesions and similar in shape to those found in uninfected brain tissues (Fig. 5A). Counts of microglia per field of cortical tissue excised from mice infected with H99, *plb1*, or Rec1 after 7 days were determined using light microscopy. Untreated (*P*□<□0.01)-, H99 (*P*□<□0.001)-, and Rec1 (*P*□<□0.01)-infected tissue exhibited on average similar and higher number of microglia per field compared to those infected with *plb1* (Fig. 5B). Then, we examined changes in the morphology of microglia during *Cn* infection and identified five different phenotypes including activated or hypertrophic (thick soma and thick ramifications), dystrophic (not-well defined soma and thin ramifications), phagocytic or amoeboid (large soma and short-retracted ramifications; ameboid or macrophage-like), ramified (cylindrical soma and long and thin ramifications), and rod-shaped (enlarged cylindrical soma and polar ramifications; Fig. 5C). The uninfected and *plb1*-infected brains have 100% ramified or resting microglia in cortical tissue (Fig. 5D). In contrast, this microglial phenotype was not visualized in cortical tissue removed from infected mice with H99 or Rec1. H99-infected mice showed 58% activated, 16% dystrophic, 24% phagocytic, and 2% rod-shaped microglia (Fig. 5D). Rec1-infected rodents demonstrated 31% activated, 46% dystrophic, 22% phagocytic, and 1% rod-shaped microglia. Our *in vivo* data demonstrate that *Cn* PLB1 is important for brain colonization and may alter microglia morphology and effector functions.

**Fig. 5.**
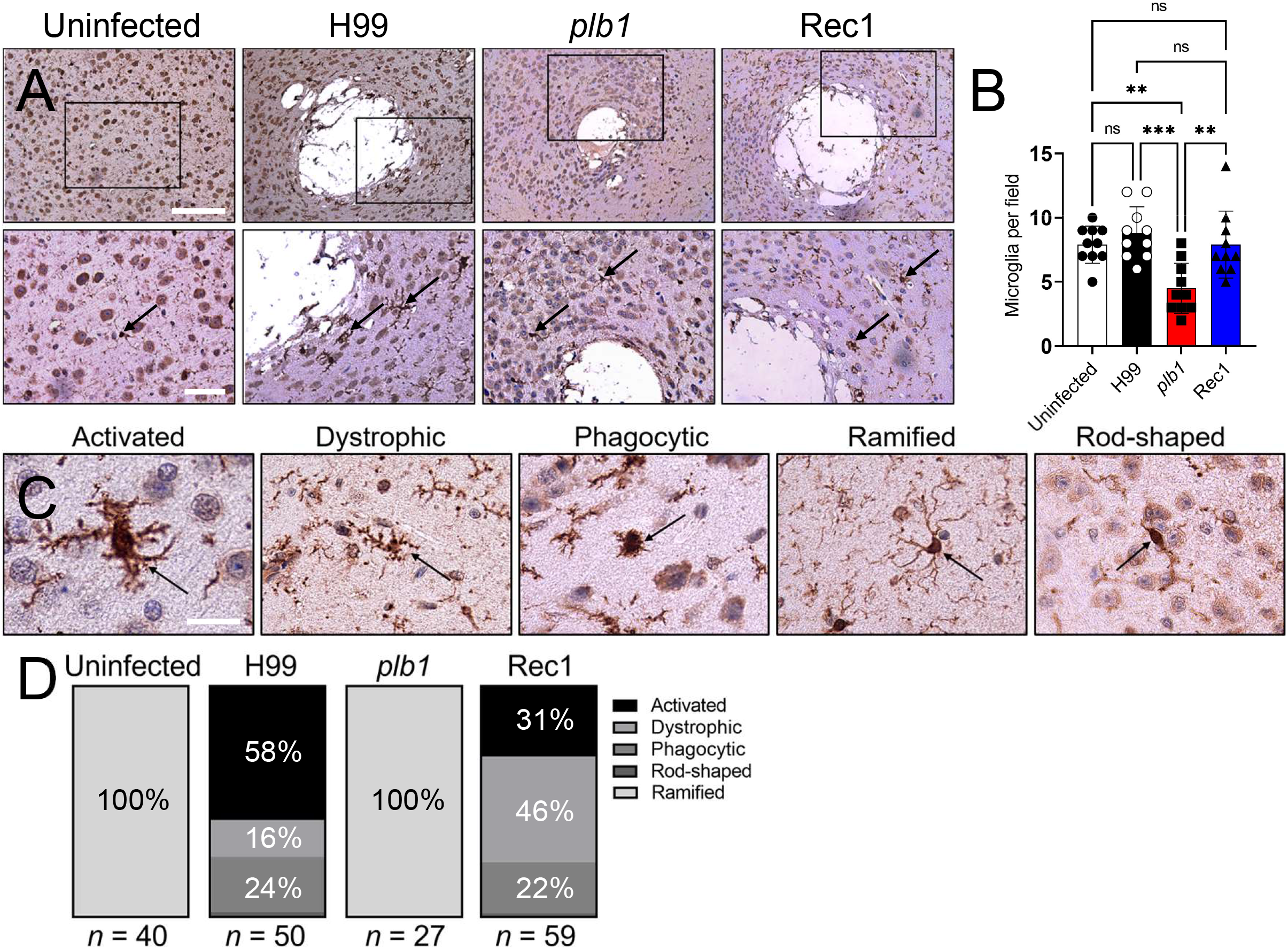
*Cn plb1* strain does not alter microglial morphology in cortical tissue. **(A)** Murine microglial responses to *Cn* strains H99, *plb1*, and Rec1 infection in cortical tissue. Uninfected mice were used as tissue baseline controls. Representative ×10 (top panel; scale bar, 100□μm), and 20 (bottom panel; scale bar, 50□μm) magnifications of ionized calcium binding adaptor protein (Iba-1)-binding mAb-stained sections of the brain are shown. Black rectangular boxes delineate the area magnified (left to right panels). Brown staining and black arrows indicate microglia. **(B)** Counts of microglia per field of cortical tissue excised from C57BL/6 mice infected with H99, *plb1*, or Rec1 after 7 days were determined using light microscopy. Bars and error bars denote the means and SDs, respectively. Each symbol represents the number of microglia per individual field (*n* = 10 per group). **(C)** The morphology of cortical microglia during cryptococcal infection consists of the following phenotypes: activated or hypertrophic, dystrophic, phagocytic or amoeboid, ramified, and rod-shaped cells. Each microglial phenotype is identified with an arrow. Scale bar, 50 μm. **(D)** Microglial type abundance during cerebral cryptococcosis. Microglia in cortical tissue of uninfected or *Cn* H99-, *plb1*-, or Rec1-infected mice were visualized under the microscope and classified according to their morphology as activated, dystrophic, phagocytic, ramified, or rod-shaped.

### *Cn plb1* strain is phagocytosed and killed by NR-9460 microglia-like cells

We investigated the efficacy of GXM-specific mAb 18B7- or complement-mediated phagocytosis of *Cn* H99, *plb1*, or Rec1 by NR-9460 microglia-like cells after 2h at 37°C and 5% CO_2_ (Fig. 6). *plb1* cells were phagocytosed more efficiently than H99 (mAb 18B7, *P*<0.001; complement, *P*<0.01) or Rec1 (mAb 18B7, *P*<0.05; complement, *P*<0.01) cells by microglia-like cells (Fig. 6A-B). Ab- and complement-mediated phagocytosis of H99 and Rec1 was similar. Then, we used two methods to determine killing of phagocytosed cryptococci by microglia, the traditional CFU (Fig. 6C) and acridine orange (Fig. 6D) assays. Significantly more PLB1 mutant cells were killed by microglia-like cells than H99 (*P*<0.0001) or Rec1 (*P*<0.0001) after 24 h co-incubation at 37°C and 5% CO_2_ (Fig. 6C). Rec1 cryptococci were killed considerably more than H99 cells (*P*<0.05). To validate the CFU killing determinations, we used acridine orange to document the percentage of cryptococcal killing in real-time as fluorescent cells (green, alive; red, dead) are directly observed under the microscope. Both, *plb1* (~76.8%; *P*<0.0001) and Rec1 (~71.3%; *P*<0.01) cells were more susceptible to killing by NR-9460 cells than H99 (~59.7%) cells (Fig. 6D). Since abrupt changes on the fungal surface negativity may influence the interactions and engulfment of cryptococci by phagocytes (24–26), we performed Zeta potential (Fig. 6E) and conductance (Fig. 6F) analyses on cells from each strain. We did not observe differences in the negative charge of H99 (−28.5±9.1 mV), *plb1* (−31.6±6.7 mV), and Rec1 (−24.5±6.8 mV) cell surfaces (Fig. 6E). However, we observed significant differences (*P*<0.0001) in cellular conductance in the following order: Rec1 (~144±1 μS) > *plb1* (~135±1 μS) > H99 (~105±0 μS). These results suggest that the lack of PLB1 activity make *Cn* prone to phagocytosis and killing by microglia, thus, impacting the ability of the fungus to colonize the CNS.

**Fig. 6.**
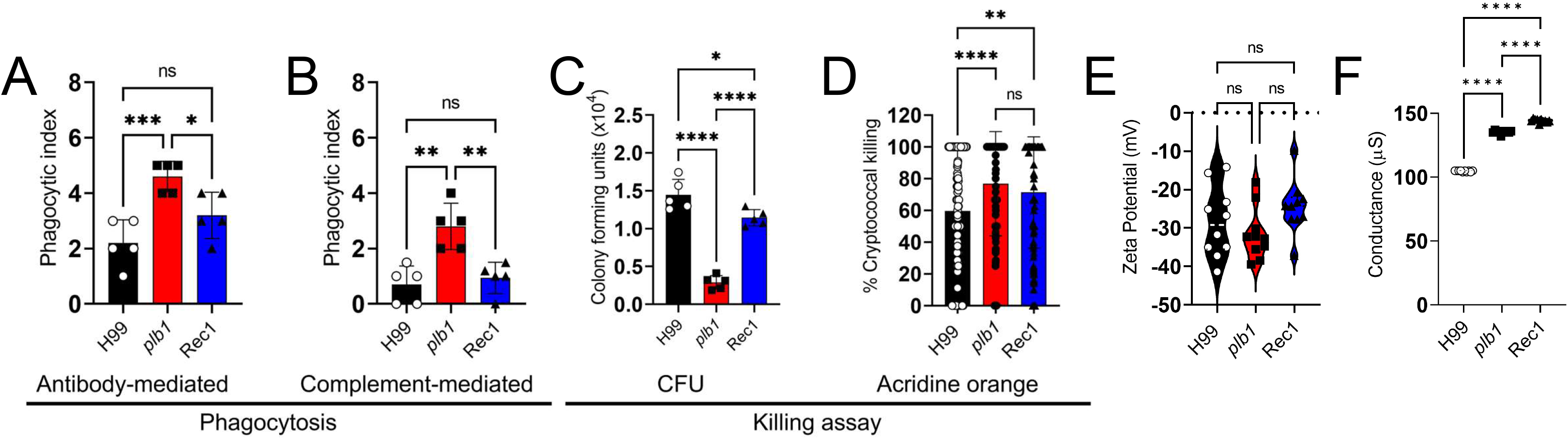
*Cn plb1* strain cells are easily phagocytosed and killed by NR-9460 microglia-like cells. The phagocytic indices (ratio of number of intracellular yeast cells to the number of microglia counted) were determined after 2 h incubation of 10^5^ NR-9460 microglia-like cells with **(A)** mAb 18B7 (IgG1)- or **(B)** complement-opsonized 10^6^ *Cn* strains H99, *plb1*, and Rec1. **(C)** CFU determinations were performed after 24 h incubation of engulfed fungi by microglia and Ab-mediated phagocytosis. **(D)** Percentage of *Cn* H99, *plb1*, and Rec1 cells killed by NR-9460 microglia-like cells. For **A** to **D**, bars represent the means of 5 independent experiments (each symbol represents 1 field consisting of 100 macrophages) and error bars indicate SDs. **(E)** Zeta potential and **(F)** conductance differences between H99, *plb1*, and Rec1 cells. Violin plot or symbols represent the means of 10 independent measurements and error bars indicate SDs. For **A** to **F**, asterisks denote *P* value significance (****, *P*□<□ 0.0001; ***, *P*□<□ 0.001; **, *P*□<□ 0.01; *, *P*□<□ 0.05) calculated by ANOVA and adjusted using Tukey’s post hoc analysis. ns denotes comparisons that are not statistically significant.

### *Cn* PLB1 mutant does not stimulate astrocytic responses in the cortex and hippocampus

Astrocytes become reactive upon interacting with *Cn* during infection (27, 28) and likely play a critical role in cryptococcal brain invasion and meningoencephalitis development. Hence, we examined under the microscope the astrocytic responses and morphological changes in murine brain tissue after a week systemic infection with *Cn* strains H99, *plb1*, or Rec1 (Fig. 7). Astrocytes in cortex, hippocampus, and cerebellum were stained with glial fibrillary acidic protein (GFAP)-binding mAb, a specific marker for astrocytes. IHC of the cortex (Fig. 7A) showed that uninfected and *plb1*-infected brains had similarly low numbers of astrocytes near to the glia limitans region of the pia matter. H99 and Rec1-infected brains displayed significantly higher astrocytosis (H99 vs. uninfected or *plb1*, *P*<0.001; Rec1 vs. uninfected or *plb1*, *P*<0.05) (Fig. 7D) and astrogliosis (H99 vs. uninfected or *plb1*, *P*<0.0001; Rec1 vs. uninfected or *plb1*, *P*<0.0001) (Fig. 7E-F) in cortical tissue than uninfected or *plb1*-infected rodents. There were no differences in astrocytosis in tissue infected with H99 or Rec1, although astrocytes in tissues infected with H99 exhibited significantly higher processes number (*P*<0.01; Fig. 7E) and thickness (*P*<0.0001; Fig. 7F) than Rec1. In the hippocampus, H99 and Rec1-infected brains displayed significantly higher astrocytosis (H99 vs. uninfected or *plb1*, *P*<0.0001; Rec1 vs. uninfected, *P*<0.0001 or *plb1*, *P*<0.001; Fig. 7B-G) and astrogliosis (H99 or Rec1 vs. uninfected or *plb1*, *P*<0.0001; Fig. 7B, H-I) compared to uninfected or *plb1*-infected rodents. No differences in astroglia number and morphology were observed between H99 and Rec1-infected hippocampi. *Cn* H99 (*P*<0.0001)-, *plb1* (*P*<0.05)-, and Rec1 (*P*<0.001)-infected cerebella displayed substantially increased astrocytosis relative to uninfected tissue (Fig. 7C-J). Likewise, H99-infected tissue had significantly higher astrocytes than *plb1*-infected tissue (*P*<0.05; Fig. 7J). Rec1 showed no differences in astrocytosis when compared to the other groups. Interestingly, Rec1 had the highest number of processes per astrocyte in the cerebellum (Fig. 7K). Both, H99 and Rec1, showed considerably higher number of processes per glia than *plb1*-infected tissue. Astrocytes interacting with H99 showed significant astrogliosis in the cerebellum and evinced the thickest processes (Fig. 7L). *plb1*- and Rec1-infected cerebella had astrocytes with similar size processes while naïve mouse brains had astroglia with the thinnest processes. These findings show that *plb1* evokes a weaker astrocytic response than H99 or Rec1 and this response depends on the brain region infected.

**Fig. 7.**
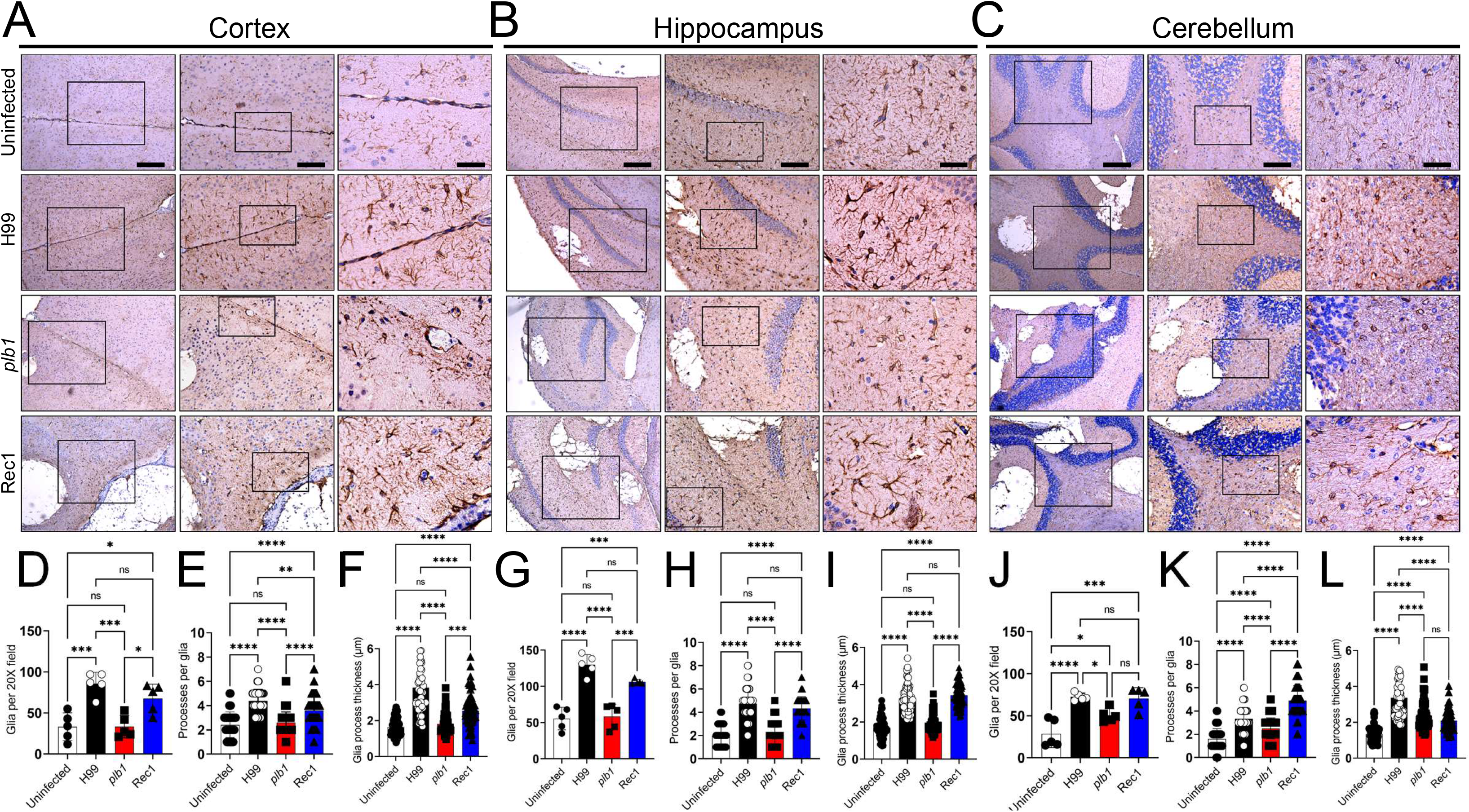
*Cn plb1* strain does not stimulate astrogliosis or morphological changes in cortical and hippocampal tissue. Histological examinations of the **(A)** cerebral cortex, **(B)** hippocampus, and **(C)** cerebellum from brains removed from *Cn* H99-, *plb1*-, or Rec1-infected mice with 10^5^ cryptococci at 7-dpi. Uninfected mice were used as tissue baseline controls. Representative ×4 (left panel; scale bar, 200□μm), 10 (center panel; scale bar, 100□μm), and 20 (right panel; scale bar, 50□μm) magnifications of glial fibrillary acid protein (GFAP)-binding mAb-stained sections of the brain are shown. Black rectangular boxes delineate the area magnified (left to right panels). Brown staining indicates astrocytes. Astrocytosis (Glia per ×20 field), processes per glia, and glia process thickness (μm) were determined in **(D-F)** cortical, **(G-I)** hippocampal, and **(J-L)** cerebellar tissue sections using an inverted microscope. For **D** to **L**, bars and error bars denote the means and SDs, respectively. Each symbol represents an individual field (*n* = 5 per group) or cell (*n* = 15 per group for processes per glia; *n* = 25 per group for process thickness). Asterisks denote *P* value significance (****, *P*□<□ 0.0001; ***, *P*□<□ 0.001; **, *P*□<□ 0.01; *, *P*□<□ 0.05) calculated by ANOVA and adjusted using Tukey’s post hoc analysis. ns denotes comparisons that are not statistically significant.

### *Cn* infection of the brain induces minimal nitric oxide (NO) production

Cryptococcal meningoencephalitis in individuals with HIV is characterized by minimal inflammation (28) while NO has anti-cryptococcal activity (29). We assessed the ability of *Cn* strains to stimulate NO production in mice and by glial-like cells. Notably, all the groups of mice showed similar accumulation of nitrite, an indicator of NO synthesis, in brain tissue (Fig. 8A). We also tested the NO production ability of NR-9460 microglia-like cells and C8-D1A, an astrocyte-like cell line isolated from mouse cerebellum, after their co-incubation without (medium, C−, negative control) or with H99, *plb1*, or Rec1 cells. Glia-like cells were incubated with lipopolysaccharide (0.5 µg/mL of LPS, component of outer membrane of Gram-negative bacteria and a potent stimulator of NO production and inflammation), interferon-γ (100 units (U)/mL of IFN-γ, a critical cytokine for immunity against microbes), and mAb 18B7 (10 µg/mL) as a positive control (C+) or with each strain. Both, NR-9460 (Fig. 8B) and C8-D1A (Fig. 8C) cells, were unable to produce NO after co-incubation with C− or with H99, *plb1*, or Rec1 for 24 h at 37 °C and 5% CO_2_. As expected, glia-like cells incubated with C+ synthesized significant amounts of NO (Fig. 8B-C). Interestingly, *Cn* inhibited NO production by NR-9460 cells after C+ was supplemented and like fungi grown with C− (Fig. 8B). In contrast, while C8-D1A cells grown with *Cn*/C+ also showed NO inhibition compared to C+, the production of the gas was higher than astrocytes grown with *Cn*/C− (*P*<0.0001; Fig. 8C). Finally, we performed western blot analysis to determine whether any *Cn* strain stimulate the expression of inducible NO synthase (iNOS) by NR-9460 and/or C8-D1A cells. We validated the results obtained *in vivo* (Fig. 8A) and *in vitro* (Fig. 8B-C) and confirmed that none of the cryptococcal strains stimulated the expression of iNOS when grown in C− conditions, thus, limiting glia-like cell NO production (Fig. 8D). NR-9460 cells incubated with H99 or *plb1* (C+) showed higher iNOS expression than C+ or Rec1 whereas C8-D1A cells displayed considerable iNOS expression when incubated in C+ or *Cn*/C+, but slightly reduced in *Cn*/C+. These results indicate that *Cn* alone does not stimulate NO production in the mouse brain or cell lines supporting the idea that cryptococcal meningitis causes minimal inflammatory response making difficult the control of the disease especially in individuals with compromised immunity. In addition, the response of astrocytes to *Cn* suggests that these cells have an important role in fighting the infection.

**Fig. 8.**
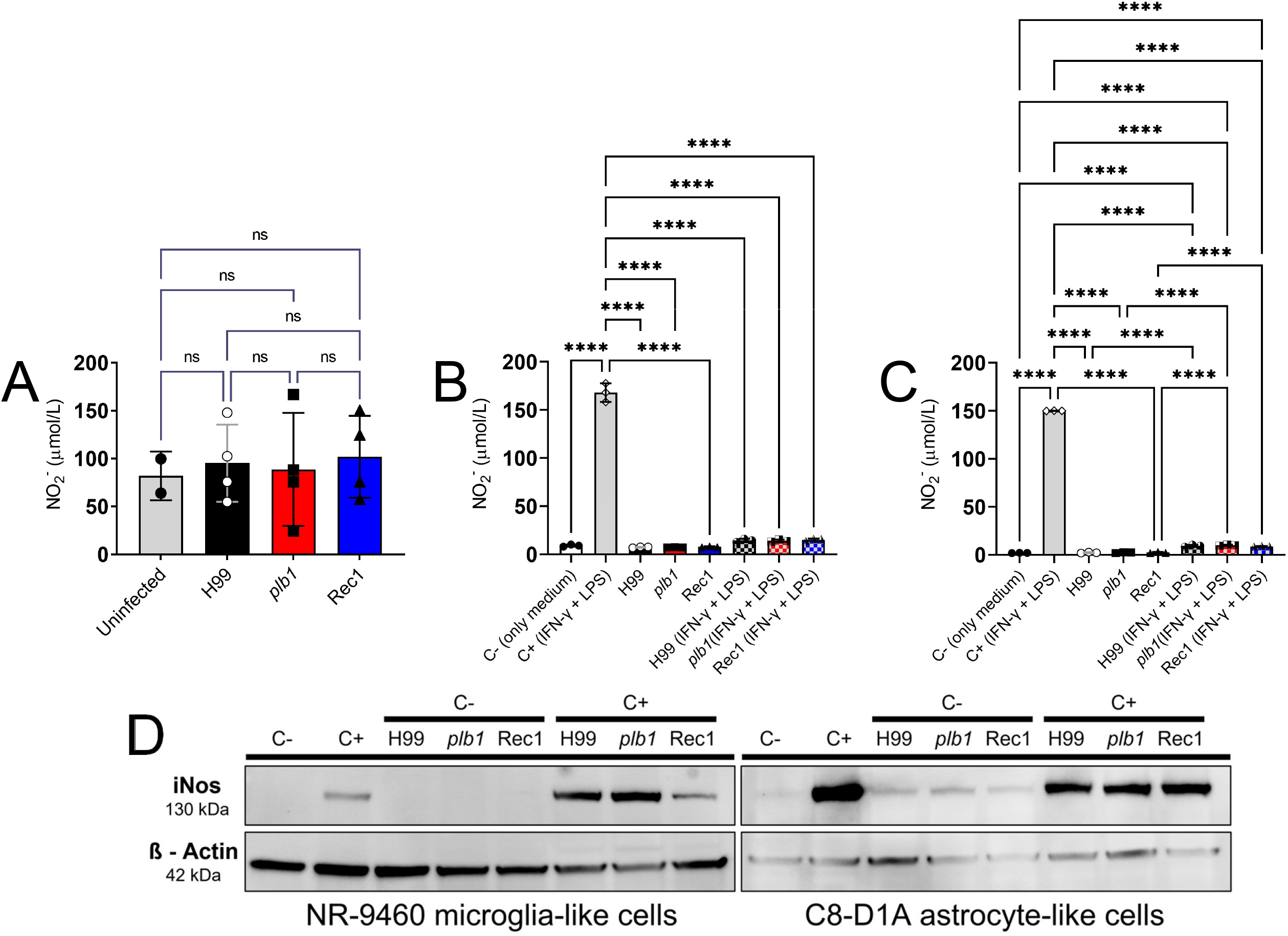
*Cn* infection of the brain induces limited nitric oxide production. **(A)** Total nitric oxide (NO_2_^−^) production was quantified based on the enzymatic conversion of nitrate to nitrite by nitrate reductase in brain tissue homogenates excised from uninfected- and H99-, *plb1*-, or Rec1-infected mice after 7 days. NO_2_^−^ production was quantified using the Griess method after **(B)** NR-9460 microglia-like and **(C)** C8-D1A astrocyte-like cells were incubated alone (control, C−, only medium) or co-incubated with interferon-γ (IFN-γ, 100 units (U)/mL), bacterial lipopolysaccharide (LPS, 0.5 μg/mL), and mAb 18B7 (10 µg/mL) alone (C+), H99 (C− or C+), *plb1* (C− or C+), Rec1 (C− or C+) for 24 h at 37 °C and 5% CO_2_. For **A** to **C**, bars and error bars denote the means and SDs, respectively. Each symbol represents an individual well (*n* = 2-4 per group). Asterisks denote *P* value significance (****, *P*□<□ 0.0001) calculated by ANOVA and adjusted using Tukey’s post hoc analysis. For **A**, ns denotes comparisons that are not statistically significant. For **B**-**C**, only comparisons with *P* value less than 0.05 are shown. **(D)** The expression of inducible NO synthase (iNOS) in NR-9460 and C8-D1A cells was determined by western blot analysis. β-actin was used as housekeeping gene control. All conditions were run in the same gel next to each other and processed in parallel. We cropped each gel to improve the clarity and conciseness of the presentation. Each gel was run three times from 3 separate protein extracts to validate the results.

## DISCUSSION

We demonstrated that *Cn* PLB1 is critical for CNS survival in mice. *plb1*-infected mice exhibited prolonged survival than those infected with H99 or Rec1. The brains of mice infected with *plb1* cryptococci exhibited comparable fungal load than those infected with H99 or Rec1 3-dpi, suggesting that *plb1* cells reach the CNS at similar rates than the other strains. However, the lack of PLB1 delays the development of the disease. A substantial difference in brain tissue fungal burden and cryptococcoma formation was observed between the *plb1* infection and the H99 or Rec1 strains 7-dpi, validating that PLB1 is essential for the fungus to survive in the CNS.

*Cn* PLB1 or manipulation of the *plb1* gene have important implications in cryptococcoma formation and CPS production/accumulation during infection. *plb1*-infected brain lesions had less GXM deposition around cryptococcomas than H99- and Rec1-infected brains. PLB1 is required for *Cn* capsule enlargement during interactions with macrophages and amoeba (14). Our SEM images confirmed that *plb1* cells also had smaller capsules compared to H99 cells. In contrast, GXM accumulation around the brain lesions in Rec1-infected mice was more extensive than in H99-infected brains. Although Rec1 cryptococci showed similar smaller capsule sizes than *plb1* cells, it is possible that the complemented strain shows smaller capsule because of enhanced CPS release in the medium. This is not the case for *plb1* cells because the histopathology demonstrated low GXM shedding around the cryptococcoma even in similar or larger size brain lesions relative to those in H99- and Rec1-infected brains. It is also conceivable that during the reconstitution of the *plb1* gene in the Rec1 strain, there were slight genomic changes that altered single cell capsule enlargement by promoting CPS hyper-production and - release. This hypothesis is supported by the biophysical data, which indicate that even though H99 and Rec1 demonstrated similar Zeta potential and complex viscosity, there were major differences in conductance, capsular fiber size, storage module, loss module, and damping behavior.

Microglia is a dynamic cell that can adapt and change its morphology according to different pathological situations (30). Surprisingly, *plb1* cortical infection shows low number of microglia and no microglial morphological changes from their ramified baseline near or around the cryptococcomas, which were indistinguishable from the naïve mice. Ramified microglia are thought to be actively “surveying” changes in their environment (31). These findings suggest that *Cn* PLB1 may be responsible for the activation and migration of microglia to the cryptococcoma, since *plb1*-infected brains had lower microglial recruitment than even uninfected brains. Alternatively, peripheral macrophages instead of microglia may be activated and recruited to control or actively participate in the infection by helping the fungus cross the BBB via the trojan horse mechanism. The perivascular immune response to H99 infection consists of monocyte, T-cell, and neutrophil infiltration, as well as substantial production of pro-inflammatory cytokines (32). Additionally, upon *Cn* intravenous infection, monocytes are primarily recruited to the brain vasculature and transport the fungus from the blood vessel lumen to the brain parenchyma (33). In contrast, *Cn* PLB1 synthesis causes microglial morphological alterations in cortical tissue. Microglia in the cortex of H99- and Rec1-infected brains showed different phenotypic abundance. Hypertrophic microglia were most abundant in H99-infected brains and are thought to be actively responding to injury (34, 35). Thus, accumulation of activated microglia may occur in damaged blood vessels or microinfarctions during cryptococcal infection. *Cn* can induce extensive fibrosis of the subarachnoid space, which may compress small veins mechanically inducing venule congestion and massive cerebral infarction (36). Brain fibrosis and infarction are associated with high mortality in HIV-cryptococcal meningitis (37). Dystrophic microglia that present fragmented or “beaded” processes, possibly due to microglial dysfunction, are significantly increased in older individuals with Down syndrome (38) and Alzheimer’s disease (39). Dystrophic microglia were present considerably in Rec1-infected brain and to a lower extent in H99-infected brains, particularly near cryptococcomas and released GXM. The accumulation of dystrophic microglia is associated with cell dysfunction and vulnerability to brain degeneration (40) or senescence (38) due to aging or neurological diseases. Rec1 released higher GXM in the cortex and hippocampus than H99, and the presence of dystrophic microglia may be related to differences in mouse behavior, which is a possibility that can be tested in future studies. Both H99 and Rec1 brains had a similar number of amoeboid microglia, which resemble macrophages due to their reduced or retracted processes. We tested the ability of NR-9460 cells to phagocytose and kill *Cn* through the Ab- and complement-mediated mechanisms and found that *plb1* was easily engulfed-killed by microglia via both mechanisms. H99 and Rec1 were similarly phagocytosed by NR-9460 cells, but Rec1 cells were more susceptible to killing than H99 cells. Even though the differences in killing between H99 and Rec1 cannot be attributed to changes in Zeta potential, Rec1 and *plb1* cells showed a similar increase in cell conductance compared to H99 cells, and the alterations in Rec1 CPS shown in the biophysical studies explain the susceptibility of this strain to killing by microglia.

Astrocytes are the most abundant glial cells in the CNS and have a vital role in neuronal homeostasis and BBB regulation. H99 and Rec1 infection promoted astrocytosis and astrogliosis compared to *plb1* infection, particularly in the cortex and hippocampus. Astrogliosis is a hallmark indicator of CNS injury, representing an early change designed to isolate and subsequently repair tissue changes independent of the underlying cause (41). The aggregation of astrocytes in the glia limitans region of the pia matter may demonstrate an anti-cryptococcal invasion mechanism. For instance, the glia limitans of the olfactory bulb is a peripheral-CNS immunological barrier to bacterial infection (42). Moreover, astrocytes can serve as antigen presenting cells in neurological (43) and autoimmune (44) diseases. *Cn* increases the expression of MHC II molecules on the surface of a human astrocyte cell line (45), suggesting that astroglia may be important to combat the fungus during CNS infection. Human astrocytes reduce *Cn* proliferation via NO-mediated mechanisms (46). Still, *Cn* can neutralize the antifungal action of NO in macrophages and astrocytes without altering the iNOS activity (47). Our results demonstrate that *Cn* impairs the ability of glia to express iNOS and NO synthesis, and together with its morphological alterations these observations support the notion that in HIV+ individuals the fungus can limit the inflammatory responses. These findings also suggest that the neurotropism shown by *Cn* in these patients is exacerbated by the inability of glia to respond to the fungus or perhaps, the emphasis of the response instead of being defensive just focuses on tissue repair after fungal-induced damage. Further studies on glia phenotypes and their function in cerebral cryptococcosis may lead to new evidence on glia-*Cn* interactions or explain the fungal neurotropism shown in immunosuppressed individuals.

## MATERIALS AND METHODS

### Cn

*Cn* isogenic strains H99, *plb1* and Rec1 were kindly provided by John Perfect at Duke University. *plb1* is a PLB1 mutant and a ura5 auxotroph of H99 with a single insertion transformed by using biolistic DNA delivery with a knockout construct containing URA5 inserted into PLB1 (16). Rec1 or reconstitution of the *plb1* strain was performed by transforming the mutant strain with a construct containing the entire *plb1* gene and the selectable antibiotic resistance gene HygB (48). *plb1* and Rec1 exhibit similar growth rate, melanin production, and capsule size to those of H99. Rec1 also shows similar phospholipase synthesis and secretion relative to the H99 (16). Yeasts were grown in sabouraud dextrose broth (pH 5.6; BD Biosciences) for 24 h at 30°C in an orbital shaker (New Brunswick Scientific) set at 150 rpm (nominal - to early stationary phase). Growth was assessed in real time at an optical density (OD) of 600□nm every 2 h using a microplate reader (Bioscreen C; Growth Curves USA).

#### Glia-like cell lines

The murine microglial cell line NR-9460 (BEI Resources, NIAID, NIH) and astrocyte cell line C8-D1A [Astrocyte type I clone] (ATCC CRL2541) are derived using brain tissue from wild-type mice. NR-9460 cells were immortalized by infection with the ecotropic transforming replication-deficient retrovirus J2. C8-D1A cells are astrocytes isolated from the cerebellum of mice. Characterization based on immunofluorescence, stimulation assays, and flow cytometry demonstrated that the NR-9460 and C8-D1A cell lines retain their glia-specific morphological, functional, and surface expression properties.

#### Systemic infection

Male and female C57BL/6 mice (6-8 weeks old; 20-25 grams; Envigo) were injected in their tail vein with a 100-μL suspension containing 10^5^ *Cn* H99, *plb1*, or Rec1 cells. Animals were monitored for survival. In separate infections, mice were also euthanized at 3-, 7-, and 14-dpi, and brain tissues were excised, photographed, weighed, and processed for determination of colony forming units (CFU; left brain hemisphere) and histopathological studies (right brain hemisphere). All animal studies were conducted according to the experimental practices and standards approved by the Institutional Animal Care and Use Committee (IACUC) at the University of Florida (protocol number: 202011067). The IACUC at the University of Florida approved this study.

#### Brain fungal load determinations

Brain tissues were homogenized in sterile phosphate-buffered saline pH 7.3 ± 0.1 (PBS). Serial dilutions of homogenates were performed; a 100-μL suspension of each sample was then plated on sabouraud dextrose agar (BD Biosciences) plates and incubated at 30°C for 48 h. Quantification of viable fungal cells was determined by CFU counts, and the results were normalized per gram of tissue.

#### Histology

The tissues were fixed in 4% paraformaldehyde (Thermo Fisher) for 24 h, processed, and embedded in paraffin. Four-micrometer coronal sections were cut, fixed onto glass slides, and subjected to hematoxylin and eosin (H&E) staining to examine cortical, hippocampal, and cerebellar tissue morphology and cryptococcoma formation. The size and number of cryptococcomas per field were determined. GXM (mAb 18B7 is an anti-cryptococcal GXM IgG1 generated and generously provided by Arturo Casadevall at Johns Hopkins Bloomberg School of Public Health; 1:1000 dilution), Iba-1 (rabbit anti-human Iba-1; 1:1000 dilution; FujiFilm Wako), and GFAP (rabbit anti-human GFAP 2033X; 1:2000 dilution; DAKO) specific Ab (conjugated to horseradish peroxidase; dilution: 1:1000; Santa Cruz Biotechnology) immunostaining to assess capsular release and distribution, microglial phenotype, and astrocyte morphology, respectively. The slides were visualized using a Leica DMi8 inverted microscope, and images were captured with a Leica DFC7000 digital camera using the LAS X digital imaging software. The number of microglia and astrocytes per region was quantified by cell counts using the recorded ×40 images and standardized 250 × 250 µm^2^ squares near cryptococcomas. The GXM distribution and number and size of processes per astrocyte in the cortex, hippocampus, and the cerebellum were calculated using the NIH Image J software (version 1.53n). Three different people blindly analyzed each parameter, and the average results are presented.

#### SEM

Cryptococci were washed three times in PBS pH 7.3 ± 0.1 and fixed in 2.5% glutaraldehyde solution grade I (Electron Microscopy Sciences) in sodium cacodylate buffer 0.1 M pH 7.2 ± 0.1 for 45 min at room temperature. Then, the cells were washed three times in 0.1 M sodium cacodylate buffer pH 7.2 ± 0.1 containing 0.2 M sucrose and 2 mM MgCl_2_ (Merck Millipore), and adhered to 12 mm diameter round glass coverslips (Paul Marienfeld GmbH & Co) previously coated with 0.01% poly-L-lysine (Sigma) for 20 min. Adhered cells were then gradually dehydrated in an ethanol growing series 30, 50 and 70% for 5 min and 95% and 100% twice for 10 min (Merck Millipore). The coverslips were then critical-point-dried using an EM DPC 300 critical point drier (Leica) and mounted on specimen stubs using a conductive carbon adhesive (Pelco Tabs™). Next, the samples were coated with a thin layer of gold or gold-palladium (10-15 nm) using the sputter method (Balzers Union). Finally, samples were visualized in a scanning electron microscope Carl Zeiss Evo LS 10 operating at 10 kV with an average working distance of 10 mm and images were collected with their respective software packages.

#### CPS biophysical analyses

*Cn* H99, *plb1*, and Rec1 cells were grown in sabouraud dextrose broth for 24 h at 37°C. A 100 μL culture suspension for each strain was transferred to a 25 mL Erlenmeyer flask with fresh minimal medium [15 mM glucose, 10 mM MgSO_4_ 7.H_2_O, 29 mM KH_2_PO_4_, 13 mM glycine and 3 µM thiamine (Sigma)] and incubated for 7 days at 37°C to stimulate capsule size growth and capsular secretion. Secreted CPSs from each cryptococcal strain cells were collected and purified by ultrafiltration using an Amicon® system with a membrane cutoff of 10 kDa (Millipore). The effective diameter and the polydispersity of the secreted polysaccharide preparations were measured in a 1 mg/mL solution (weight per volume) by quasi-elastic light scattering in a NanoBrook Omni particle-size analyzer (Brookhaven Instruments Corp., NY). The Zeta potential (ζ) of capsular secreted polysaccharide samples were calculated on a Zeta potential analyzer (NanoBrook Omni particle). The passive micro rheology of a concentrated solution from polysaccharides released in culture from each cryptococcal strain was characterized by measuring the complex viscosity η(ω), viscous modulus (G”) and elastic modulus (G’) of these solutions in a NanoBrook Omni particle-size analyzer. Measurements were performed in the angular frequency (ω) range of 10-10^6^ rad/s. The viscoelastic behavior (tanδ), calculated as tanδ = G′′/G′, was studied using 1 μm diameter polystyrene beads (Polybead®; Polysciences) as a standard for all the experiments, which were performed in triplicate.

#### Phagocytosis assay

Monolayers of NR-9640 cells were washed thrice with PBS, and feeding medium (49, 50) supplemented with IFN-γ (100□U/mL) and LPS (0.5□µg/mL) was added, followed by the addition of preincubated cryptococci with mAb 18B7 (10□μg/mL) or complement (mouse serum) for 1□h, in a microglia (10^5^ cells):yeast (10^6^ cells) ratio of 1:10. The plates were incubated for 2□h at 37°C and 5% CO_2_ for phagocytosis. For microscopic determination of phagocytosis, the monolayer coculture was washed thrice with PBS to remove nonadherent cells, Giemsa stained, fixed with cold methanol, and viewed with light microscopy as described. The phagocytic index was determined to be the number of internalized yeast cells over the number of 100 microglia per well. Internalized cells were differentiated from attached cells because microglia with internalized cryptococci tend to grow their cell body in the direction in which the intracellular fungal cells are present.

#### Traditional killing assay

After Ab-mediated phagocytosis, each well containing interacting microglia-crytptococci was gently washed with feeding medium three times to get rid of fungal cells that were not phagocytized. Then, cryptococcus-engulfed microglia were incubated for 24□h at 37°C and 5% CO_2_. Microglia-like cells were lysed by forcibly pulling the culture through a 27-gauge needle five to seven times. A volume of 100 µL of suspension containing cryptococci was aspirated from the wells and transferred to a microcentrifuge tube with 900 µL of PBS. For each well, serial dilutions were performed and plated in triplicate onto sabouraud dextrose agar plates, which were incubated at 30°C for 48□h. Viable cryptococcal cells were quantified as CFU. Although it is plausible that noninternalized cryptococci could replicate for several generations in feeding medium during a 24-h period, wells for each condition were microscopically monitored after phagocytosis to reduce the possibility of obtaining confounding results.

#### Acridine Orange killing assay

The viability and non-viability of fungi can be differentiated during phagocytosis by acridine orange staining, while extracellular organisms are quenched with crystal violet (51). The phagocytosis of cryptococci by NR-9640 cells was performed under similar conditions as described above in an 8-chamber polystyrene tissue culture glass slide (BD Biosciences). After phagocytosis and 24 h incubation at 37°C, microglia-like cells with intracellular cryptococci were washed with Hank’s Balanced Salt Solution (pH 7.2; Thermo Fisher), and the slides were stained with 0.01% acridine orange (Sigma) for 45 sec by the method of Pruzanski and Saito (52). The slides were gently washed with Hank’s Balanced Salt Solution and stained for 45 sec with 0.05% crystal violet (Sigma) dissolved in 0.15 M NaCl (Sigma). Finally, the slides were rinsed 3 times with PBS, mounted on microscope coverslips, and sealed at the edge with nail polish. The percentage of fungal-related cell killing was determined by fluorescence microscope (Leica DMi8 inverted microscope). Intracellular living and dead cryptococci fluoresce green and red, respectively. For each experiment 10 fields in each well were counted per well, and at least 100 macrophages with phagocytized fungal cells were analyzed in each well.

#### NO production

Nitrite produced in the brain homogenate supernatants of *Cn* infected C57BL/6 mice and supernatants of 10^5^ NR-9460 or C8-D1A cells were quantified 24□h after incubation with 10^6^ *Cn* H99, *plb1*, or Rec1 cells alone or with 100 U/mL of IFN-γ, 0.5 µg/mL of LPS, and 10 µg/mL of mAb 18B7 in feeding medium using the Total NO and Nitrate/Nitrite Parameter (R&D Systems; detection limit, 0.78 μmol/L) and Griess Reagent method (Invitrogen; detection limit, 100 nmol/L) kits, respectively, according to the manufacturers’ protocol. NO levels were monitored by measuring the optical density at 540-548□nm using a microtiter plate reader (Bio-Tek). Feeding medium alone (C−; negative) or with LPS and IFN-γ (C+; positive) were used as controls.

#### Western blot analysis

Western blot analysis was conducted using cytoplasmic extracts from mouse brain cells made with a NE-PER nuclear and cytoplasmic extraction kit (Thermo Fisher). The mixture was centrifuged at 10,000□×□g for 10 min at 4 °C, and the resulting protein content of the supernatant was determined using the Bradford method, employing a Pierce BCA protein assay kit (Thermo Fisher). Lysates were preserved in a protease inhibitor cocktail (Thermo Fisher) and stored at □−□20 °C until use. Extracts were diluted with 2□×□Laemmli sample buffer (Bio-Rad) and β-mercaptoethanol (Sigma). The mixture was heated to 90 °C for 5 min. Twenty-three µg of protein were applied to each lane of a gradient gel (7.5%; Bio-Rad). Proteins were separated by electrophoresis at a constant 130 V/gel for 90 min and transferred to a nitrocellulose membrane on the Trans-Blot Turbo Transfer System (Bio-Rad) at 25 V for 7 min. The membranes were blocked with 5% BSA in tris-buffered saline (TBST; 0.1% Tween 20) for 2 h at RT. A primary monoclonal iNOS-specific Ab (rat anti-mouse, dilution, 1:200; Santa Cruz Biotechnology) was incubated overnight at 4 °C with TBST (5% BSA). After washing the membranes 3X with TBST for 10 min, a rabbit anti-rat conjugated to horseradish peroxidase was used as a secondary Ab (1:1000; Southern Biotech) and incubated with TBST (5% BSA) for 40 min at RT. The membranes were washed as described above. Protein bands were measured using the UVP ChemStudio imaging system (Analytik Jena) after staining each membrane with chemiluminescence detection reagents (Thermo Fisher). Housekeeping protein β-actin (a cytoskeleton protein; dilution, 1:5,000; BD Biosciences) was used as loading controls. This western blot protocol was previously described in (53) and modified accordingly for this study.

#### Statistical analysis

All data were subjected to statistical analysis using Prism 9.4 (GraphPad Software). Differences in survival rates were analyzed by the log rank (Mantel-Cox) test. *P* values for multiple comparisons were calculated by analysis of variance (ANOVA) and adjusted using the Tukey’s multiple comparison test. *P* values of □<□0.05 were considered significant.

## ACKNOWLEDGEMENTS

M.F.H., M.E.M., M.R-G, and L.R.M. were supported by the National Institute of Allergy and Infectious Diseases (NIAID award # R01AI145559) of the US National Institutes of Health (NIH). The following reagent was obtained through BEI Resources, NIAID, NIH: Microglial Cell Line Derived from Wild Type Mice, NR-9460. G.R.D.S.A. and S.F. were supported by Fundação Carlos Chagas Filho de Amparo à Pesquisa do Estado do Rio de Janeiro.

## AUTHORSHIP CONTRIBUTIONS

All authors contributed to the project design and experimental procedures, analyzed data, provided the figure presentation, and manuscript writing.

## Notes

**CONFLICT OF INTEREST**: The authors declare no conflict of interest.

### Competing Interest Statement

The authors have declared no competing interest.

